# OmicsX: a web server for integrated OMICS analysis

**DOI:** 10.1101/755918

**Authors:** Jianbo Pan, Jiang Qian, Hui Zhang

**Affiliations:** Department of Pathology, Johns Hopkins University, School of Medicine, Baltimore, MD 21287, USA; Department of Ophthalmology, Johns Hopkins University, School of Medicine, Baltimore, MD 21287, USA

## Abstract

High-throughput omic data sets like genomics, transcriptomics, proteomics, glycomics, lipitomics, metabolomics, and modified forms of the biological molecules have been generated to investigate biological mechanisms. Accordingly, bioinformatic methods and tools have been developed and applied to analyze the omics data for different purposes. However, a lack of comparison and integration tools impedes the deep exploration of multi-omics data. OmicsX is a user-friendly web server for integration and comparison of different omic datasets with optional sample annotation information. The tool includes modules for gene-wise correlation, sample-wise correlation, subtype clustering, and differential expression. OmicsX provides researchers the analysis results with visualization, searching, and data downloading, and help to suggest the biological indications from the comparison of different omic data.

**Availability and implementation:** http://bioinfo.wilmer.jhu.edu/OmicsX/

## Introduction

Omics refers to the collective analysis that measure some characteristic of a large family of cellular molecules to explore the roles of the various types of molecules that make up the cells and tissues of an organism. Omics technologies were widely and rapidly developed for universal detection of genes and their modifications (genomics, epigenomics), mRNA (transcriptomics), proteins and their modifications (proteomics and post-translational proteomics), lipids (lipidomics), and metabolites (metabolomics). The integration of these omics by systems biology has found applications in different areas, including the Cancer Proteomic Tumor Analysis Consortium (CPTAC) proteogenomic studies (Zhang,H., Liu,T., et al. 2016). Due to the decrease of time- and cost-consuming in omics techniques, researchers have generated extensive omics data. However, the interpretation of those data is a challenge.

Traditionally, clustering, principal component analysis (PCA), and differential analysis were used to analyze a specific omics data with the focus of delineating sample heterogeneity and identifying molecular changes associated with phenotypes. Accordingly, some tools like ClustVis (Metsalu,T. and Vilo,J. 2015), PANDA-view (Chang,C., Xu,K., et al. 2018), START App (Nelson,J.W., Sklenar,J., et al. 2017) and iDEP (Ge,S.X., Son,E.W., et al. 2018) have been developed. Recently, CPTAC introduced integrated proteogenomics approaches to link proteomics and genomics/transcriptomics profiles in cancer studies, utilizing analytical methods like gene expression correlation, and clustering comparisons of mRNA and protein expression profiles. Overall, as multi-omic studies become more widespread, there is a need to develop bioinformatic platforms that integrate multi-omic datasets.

Although less tools focused on the integration of multi-omics, 3Omics is a web application that allows users to input multi omics datasets to perform network analysis, correlation analysis, co-expression profiling, phenotype mapping, pathway and GO enrichment analysis (Kuo,T.C., Tian,T.F., et al. 2013). However, currently there are no tools that have sufficient flexibility to statistically compare expression datasets and associate the concordance between them with biological or other sample annotations. Here, we introduce the easy-to-use web server, OmicsX, for integrating multiple omics datasets with optional sample annotation information. Example comparative datasets examined can include before/after drug treatment profiles, transcriptomic/proteomic profiles, and cross-species profiles. The tool modules includes gene-wise correlation, sample-wise correlation, sample clustering, and differential expression. OmicsX features include data searching and result matrix downloading at both the gene level and pathway level, and visualization of select analysis results. Overall, OmicsX is a simple, bioinformatic platform that can aid researchers inferring the biological insight from the comparison omics data.

## Results

### Access of OmicsX

The OmicsX is available online at http://bioinfo.wilmer.jhu.edu/OmicsX/. Python packages were implemented to process and analyse the user-uploaded omics data. A user-friendly web interface designed using JSP technology on Ubuntu Linux operating system was provided to facilitate data searching, downloading and updating. The main pages of OmicsX included data upload page (home), download page and help page. The download page will provide download of the python packages for users to process the data in-house. And the help page will describe the parameters and figures involved in OmicsX. In the following, we briefly describe data input and four result sections of OmicsX (Fig. 1).

**Fig. 1.**
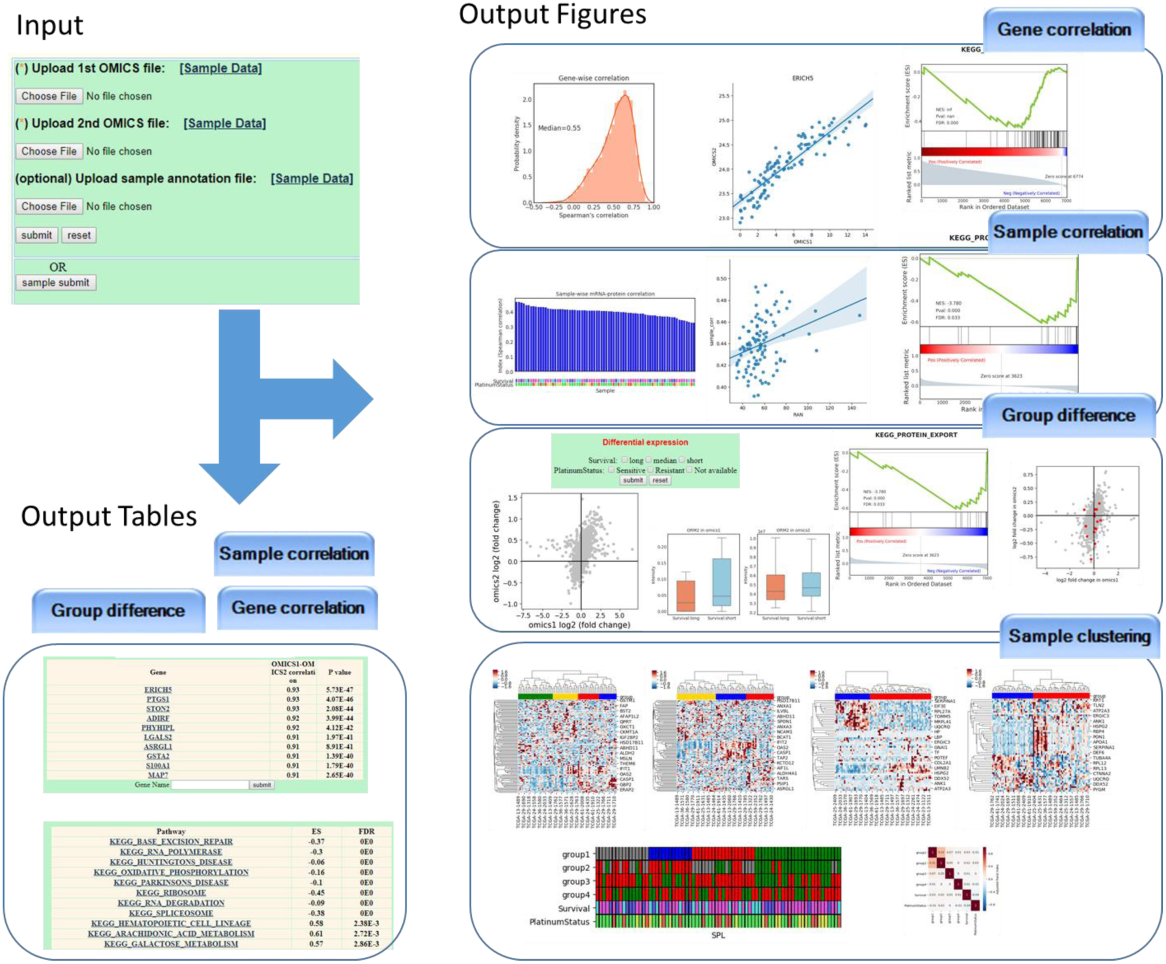
Input and output of OmicsX.

### Data Input

Normalized omics data matrix are required to initiate the analysis in the home page. The matrix should be numeric with sample names in first row and gene names in first column like commonly used format. Optionally, a sample annotation file could also be uploaded to provide the classification information for association with the integration analysis. Submission with sample data from CPTAC ovarian study (Zhang,H., Liu,T., et al. 2016) is also provided for users to familiarize themselves with the usage of OmicsX.

### Gene-wise correlation

The data matrices were firstly checked and filtered by overlapped genes and samples for further downstream integration and comparison analysis. For each overlapped gene, Spearman’s correlation is calculated across overlapped samples. The results are shown in the table and also downloadable in the result page from OmicsX. A correlation histogram is provided to visualize the consistency between two omics data from the gene-wise dimension (Fig. 2). The correlation ranked gene list is further input into Gene Set Enrichment Analysis (GSEA) for pathway enrichment analysis using GSEApy (version 0.7.0) (https://github.com/BioNinja/gseapy), a Python implementation of GSEA algorithm (Subramanian,A., Tamayo,P., et al. 2005). The significantly highly and lowly correlated pathways were listed in table. Clicking gene name and pathway name could lead to scatter plot of gene expression between two omics data and GSEA plot of significantly correlated pathway (Fig. 2).

**Fig. 2.**
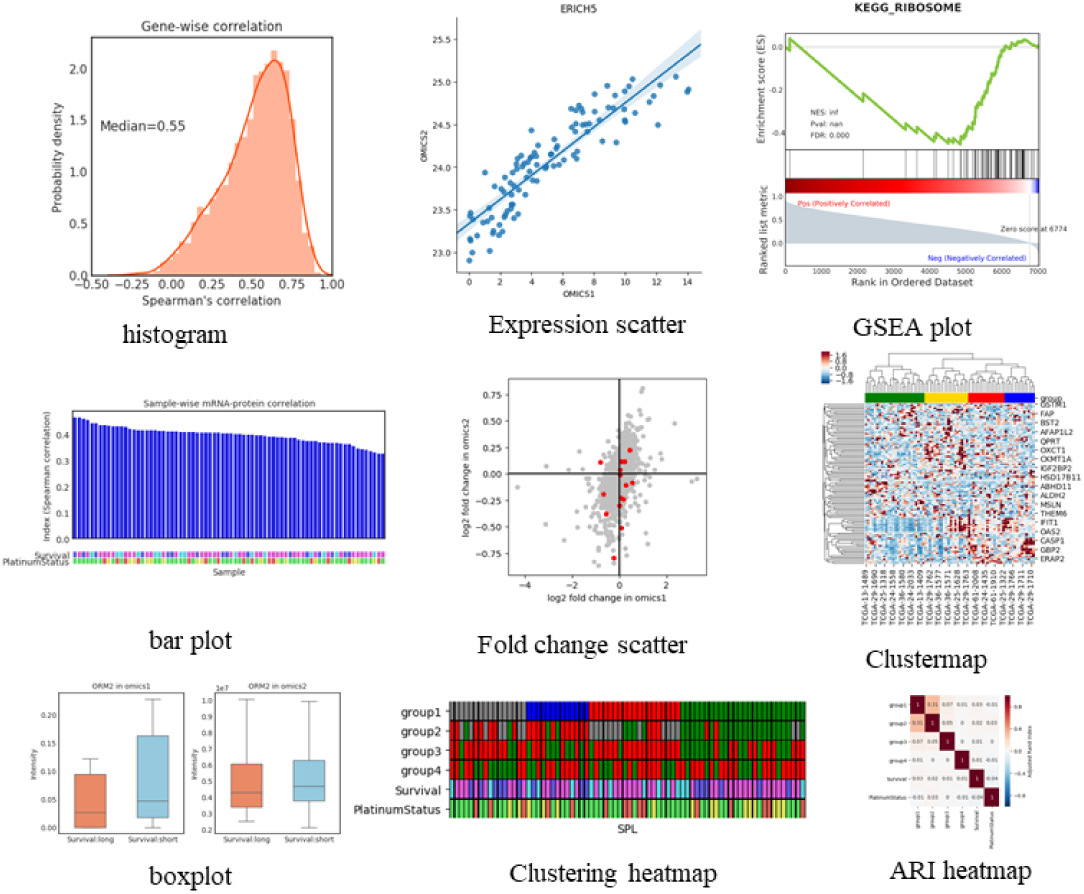
Representative output figures of OmicsX.

### Sample-wise correlation

For individual sample, sample-wise Spearman’s correlation is also calculated across overlapped genes and set as sample index for further analysis. If the sample annotation information is provided, the t test of sample index between one group and left groups will be done group by group. If any test show significance, the significance will be labelled with * in the bar plot of sample index (Fig. 2). To identify genes associated with the sample phenotypes, we calculated correlation between the sample phenotypes and gene expressions across different samples. The ranked list is also input for pathway enrichment analysis as described in the gene-wise correlation section. Similar as those in gene-wise section, tables and figures for genes and significant pathways are also provided.

### Differential analysis

Differential analysis module is available when sample annotation file is provided by the user. To start differential analysis, two annotation groups must be selected firstly. For each gene, fold change will be calculated between two groups in the first omics and the second omics data, separately. Scatter plot will be draw to show the fold change correlation between two dataset. For individual gene, the boxplot is provided (Fig. 2). By the absolute value of log2 fold change, genes are also ranked for GSEA analysis to find the consistent/different differential pathways between two omics dataset. For each pathway, enrichment plot and scatter plot are provided (Fig. 2).

### Sample Clustering

Sample heterogeneity is investigated using clustering analysis in this section. Four types of clustering are construct based on four different gene sets: the first using top half correlated genes in the first omics dataset; the second using top half correlated genes in the second omics dataset; the third using bottom half correlated genes in the first omics dataset; the fourth using bottom half correlated genes in the second omics dataset. The above four different group information and annotation information of the samples were gathered in a clustering heatmap, and adjusted Rand index (ARI) (Hubert, L. and Arabie, P. 1985) is calculated between different groups and displayed in ARI heatmap (Fig. 2). ARI, a similarity measure between two clusters, has a value close to 0.0 for random labelling independently of the number of clusters and samples and exactly 1.0 when the clusters are identical.

### Usage example

In the usage example, we initiated the analysis using RNAseq-based transcriptomics data, mass spectrometry-based proteomics data and clinical phenotype of 77 ovarian cancer tissues in CPTAC ovarian study (Zhang,H., Liu,T., et al. 2016). As shown in fig.1, a median Spearman’s correlation value of 0.55 for each mRNA-protein was observed in section of “Gene-wise correlation”. Genes with higher correlations were involved in some metabolic pathways like butanoate metabolism. In section of “Sample-wise correlation”, a median Spearman’s correlation value of 0.38 for sample-wise mRNA-protein was observed. Pathway of RNA degradation showed a significant negative association with sample-wise mRNA-protein correlations, and further indicated a possible transcription regulation in different tissues. In section of “Differential analysis”, group “long survival” and group “short survival” were selected for further analysis. The fold changes of genes between these two groups in RNA and protein data show a correlation of 0.46, and genes in pathway of aminoacyl-tRNA biosynthesis show a good consistence. In section of “Sample Clustering”, the two clustering results show a high concordance (ARI=0.31) when using those highly mRNA-protein correlated genes, and may provide a more robust subtype information.

## Conclusion

In summary, we have established a user-friendly web-server OmicsX that facilitates the comparison analysis of multiple omics data. This will enable scientific researchers, as well as, bioinformatics enthusiasts with little programming background, to be able to explore unique biology from omics data. To our knowledge, it is the most comprehensive analysis platform for examining and comparison multi-omics data, and will serve as an effective tool to aid users with omics data analysis.

## Acknowledgement

This work was supported by the National Cancer Institute, the Clinical Proteomic Tumor Analysis Consortium (CPTAC, Grant U24CA210985) and the Early Detection Research Network (EDRN, U01CA152813). The authors would like to thank Dr. David J. Clark for his assistance with reading the manuscript and other CPTAC investigators for their helpful discussions.

